# A multi-omic approach reveals iron availability influences cell fate fidelity

**DOI:** 10.1101/2025.03.19.644252

**Authors:** Athena Jessica S Ong, Tara A Tigani, Jordyn M Reinecke, Andrew G Cox, Kristin K Brown

**Author notes:** Equal contribution (Andrew G Cox, Kristin K Brown).

## Abstract

Recent evidence has highlighted the importance of employing culture media designed to emulate the metabolic environment Here, we utilize the physiological medium Plasmax to examine the impact of nutrient availability on the human hepatocyte cell line, HepG2. Incubation of HepG2 cells in Plasmax suppressed a transcriptional program driven by Hepatocyte Nuclear Factor 4 (HNF4A), a master regulator of hepatocyte identity. Given that HepG2 cells were originally isolated from a patient with hepatoblastoma, this suggests reversion to the native state in physiological media. Importantly, exclusion of iron from Plasmax reinstated the HNF4A-driven transcriptional program. These studies suggest a relationship between iron availability and the fidelity of hepatocyte cell fate and highlight the importance of more faithfully recapitulating *in vivo* metabolite availability *in vitro*.

## Introduction

*In vitro* cell culture models play a central role in biological and biomedical research. In recent years, efforts to improve the physiological relevance of these models have highlighted the importance of media composition on cellular phenotype and biological responses. For example, the dependence of some cancer cell lines on glutaminolysis is not observed in the *in vivo* context (Davidson *et al*, 2016). This has been attributed to the presence of high cystine concentrations in conventional *in vitro* culture conditions (Muir *et al*, 2017). More generally, conventional media (e.g. Dulbecco’s Modified Eagle’s Medium, DMEM) poorly recapitulate metabolite levels that cells would be exposed to *in vivo* (Eagle, 1959; Dulbecco & Freeman, 1959). The increasing awareness and recognition that the composition of cell culture media is important in preserving biological processes *in vitro* has led to ongoing efforts to develop cell culture media that better represent physiological metabolite concentrations (Vande Voorde *et al*, 2019; Cantor *et al*, 2017; Tardito *et al*, 2015). Studies employing physiological media have demonstrated that media composition can alter cellular dependencies via metabolic and transcriptional changes (Flickinger *et al*, 2024; Vande Voorde *et al*, 2019; Cantor *et al*, 2017; Apiz Saab *et al*, 2023; Rawat *et al*, 2024; Senkowski *et al*, 2023).

Hepatocytes, the major cell type in the liver, are widely modelled *in vitro* to study drug metabolism, drug-induced liver injury (DILI), and various liver diseases (Kaur *et al*, 2023; Yang *et al*, 2023). The most commonly used cell line in these contexts are HepG2 cells, which were originally derived from a hepatic tumour isolated from a 15-year-old male (Aden *et al*, 1979; Arzumanian *et al*, 2021). Although initially characterised as being derived from a hepatocellular carcinoma (HCC), subsequent histopathologic, genetic and molecular analysis revealed that the origin of HepG2 cells was instead from a hepatoblastoma (HB) (López-Terrada *et al*, 2009). Despite this, HepG2 cells are still widely utilized as a hepatocyte cell line as they exhibit the hallmarks of hepatocyte fate and function *in vitro* (Liu *et al*, 2022, 2021; Lin *et al*, 2016; Park *et al*, 2018; Zhang *et al*, 2018).

In this study, we examined the impact of nutrient availability on HepG2 cells by comparing culture in conventional media with culture in a media previously developed to more faithfully reflect metabolite abundance in human plasma (Plasmax) (Vande Voorde *et al*, 2019). We demonstrate that culturing HepG2 cells in Plasmax alters cell fate in a manner associated with loss of HNF4A. Moreover, we demonstrate that iron availability contributes to the regulation of hepatic cell fate. This research suggests an important role for physiological media in maintaining cell fate *in vitro*.

## Results

### HepG2 cells adopt a native hepatoblast-like state in physiological media

Typically, HepG2 cells are maintained in one of three conventional cell culture media: EMEM, high glucose DMEM (HG-DMEM) or low glucose DMEM (LG-DMEM) (Sanghvi *et al*, 2019; Chen *et al*, 2024; Jiang *et al*, 2023; Ong *et al*, 2023). To examine the impact of culture media on the transcriptional landscape, HepG2 cells were cultured in each of the three conventional media types or in the physiological medium Plasmax (Vande Voorde *et al*, 2019). Plasmax mimics metabolite concentrations found in human plasma and has a dramatically different composition than conventional cell culture media (Fig. S1A). While minimal transcriptional changes were observed in response to culture in any of the three conventional media, culture in Plasmax induced widespread transcriptional changes (Fig. S1B,C). Amongst the most significant changes observed in Plasmax, relative to EMEM, was upregulation of the pioneer transcription factor SRY-Box Transcription Factor 4 (*SOX4*), a regulator of hepatocyte dedifferentiation, and concomitant downregulation of genes (Apolipoprotein H, *APOH*; Albumin, *ALB*; Transferrin, *TF*) controlled by Hepatocyte Nuclear Factor 4 Alpha (HNF4A), a transcription factor that regulates hepatocyte identity (Fig. 1A) (Parviz *et al*, 2003; Sekiya & Suzuki, 2011; Katsuda *et al*, 2024). Downregulation of HNF4A in Plasmax was confirmed at the level of protein expression (Fig. 1B). Consistent with these observations, Gene Set Enrichment Analysis (GSEA) revealed downregulation of signatures associated with hepatocyte cell state and hepatocyte function in the context of culture in Plasmax (Fig. 1C). In the developmental context, hepatoblasts are bipotential progenitor cells that differentiate to give rise to both hepatocytes and cholangiocytes (Fig. S1D) (Gordillo *et al*, 2015). Culture in Plasmax induced an increase in the expression of genes associated with a hepatoblast-like state (e.g. Epithelial Cell Adhesion Molecule, *EPCAM* and SRY-Box Transcription Factor 9, *SOX9*) and, as previously mentioned, loss of genes associated with differentiated hepatocytes (e.g. *ALB, APOH*, and *TF*) suggesting that culture in physiological media drives dedifferentiation (Fig. 1D). In contrast, culture of the HCC-derived hepatocyte cell line Hep3B in Plasmax did not impact HNF4A protein expression (Fig, S1E). This was expected as Hep3B cells are not derived from HB and represent an intrinsically differentiated hepatocyte cell line. Finally, analysis of a publicly available dataset examining the impact of culture conditions on the basal-like breast cancer cell line MDA-MB-468 revealed a shift to a progenitor-like state when cells were cultured in Plasmax (Fig. S1F) (Vande Voorde *et al*, 2019; Charafe-Jauffret *et al*, 2006; Neve *et al*, 2006). Overall, these data suggest that culture in physiologic media maintains cell fate fidelity *in vitro*.

**Figure 1.**
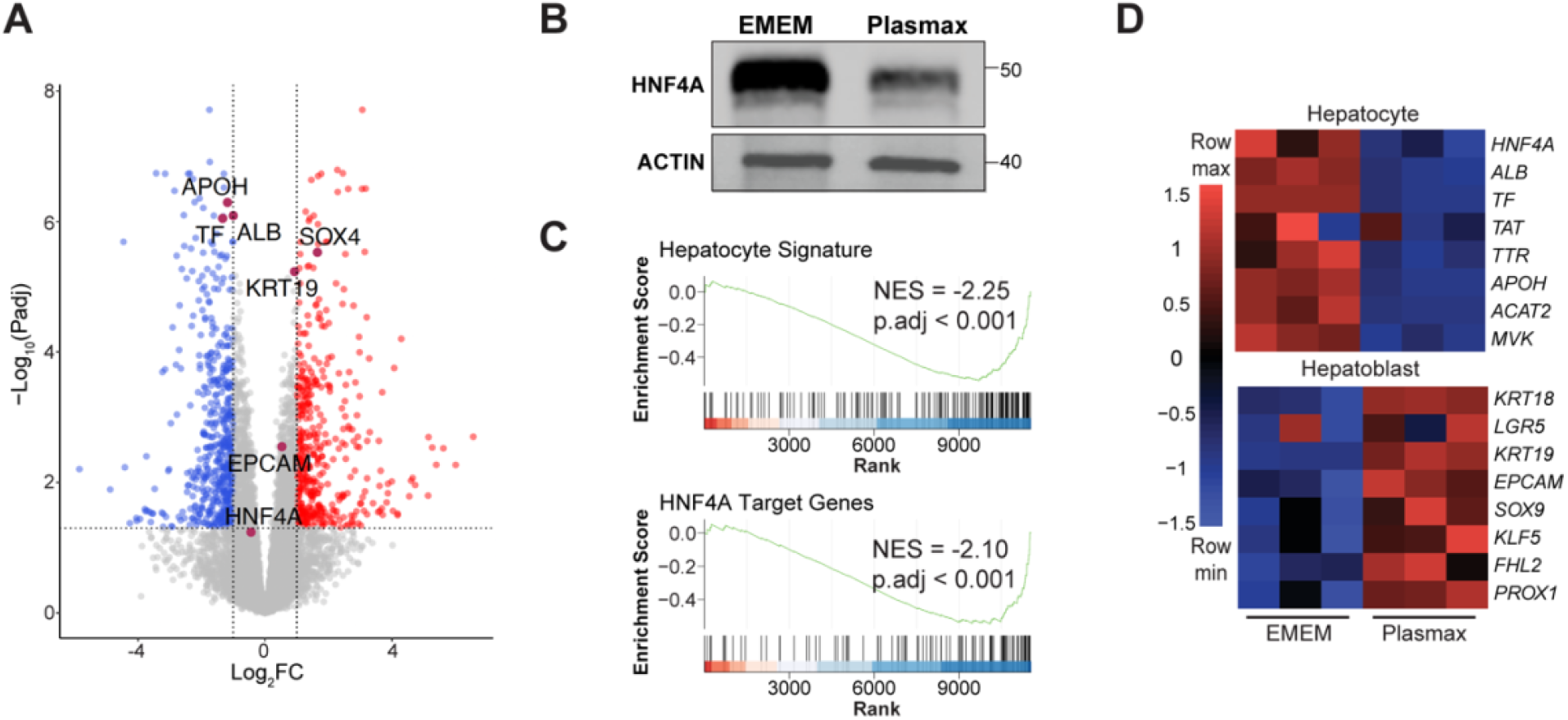
HepG2 cells adopt a native hepatoblast-like state in physiological media. (A) Volcano plot of differentially expressed genes (DEGs) identified from RNA-Seq analysis comparing HepG2 cells cultured in Plasmax with HepG2 cells cultured in EMEM, n=3. Hepatocyte differentiation markers are highlighted in maroon. Significantly upregulated genes are highlighted in red. Significantly downregulated genes are highlighted in blue. (B) Representative immunoblot analysis of HepG2 cells cultured in EMEM or Plasmax. (C) Gene set enrichment analysis (GSEA) plots derived from RNA-Seq analysis comparing HepG2 cells cultured in Plasmax with HepG2 cells cultured in EMEM demonstrating signatures associated with hepatocyte cell state and function. (D) Heatmap of hepatocyte and hepatoblast gene expression in HepG2 cells culture in EMEM or Plasmax as determined by RNA-Seq analysis, n=3.

### Trace element availability regulates hepatocyte differentiation

To examine the components of Plasmax contributing to dedifferentiation of HepG2 cells, variants of Plasmax lacking either trace elements, which are severely underrepresented in EMEM/DMEM, or lacking metabolites absent from EMEM/DMEM (termed “supplemental metabolites”) were prepared (Fig. S1A) (Vande Voorde *et al*, 2019). Withdrawal of trace elements, but not supplemental metabolites, restored HNF4A protein expression to levels observed in EMEM (Fig. 2A). Moreover, transcriptomics revealed that trace element withdrawal restored hepatocyte gene expression and expression of signatures associated with hepatocyte cell state and hepatocyte function (Fig. 2B,C). Notably, *SOX4* expression was suppressed and returned to levels observed in EMEM when cells were cultured in Plasmax devoid of trace elements (Fig. 2C,D).

**Figure 2.**
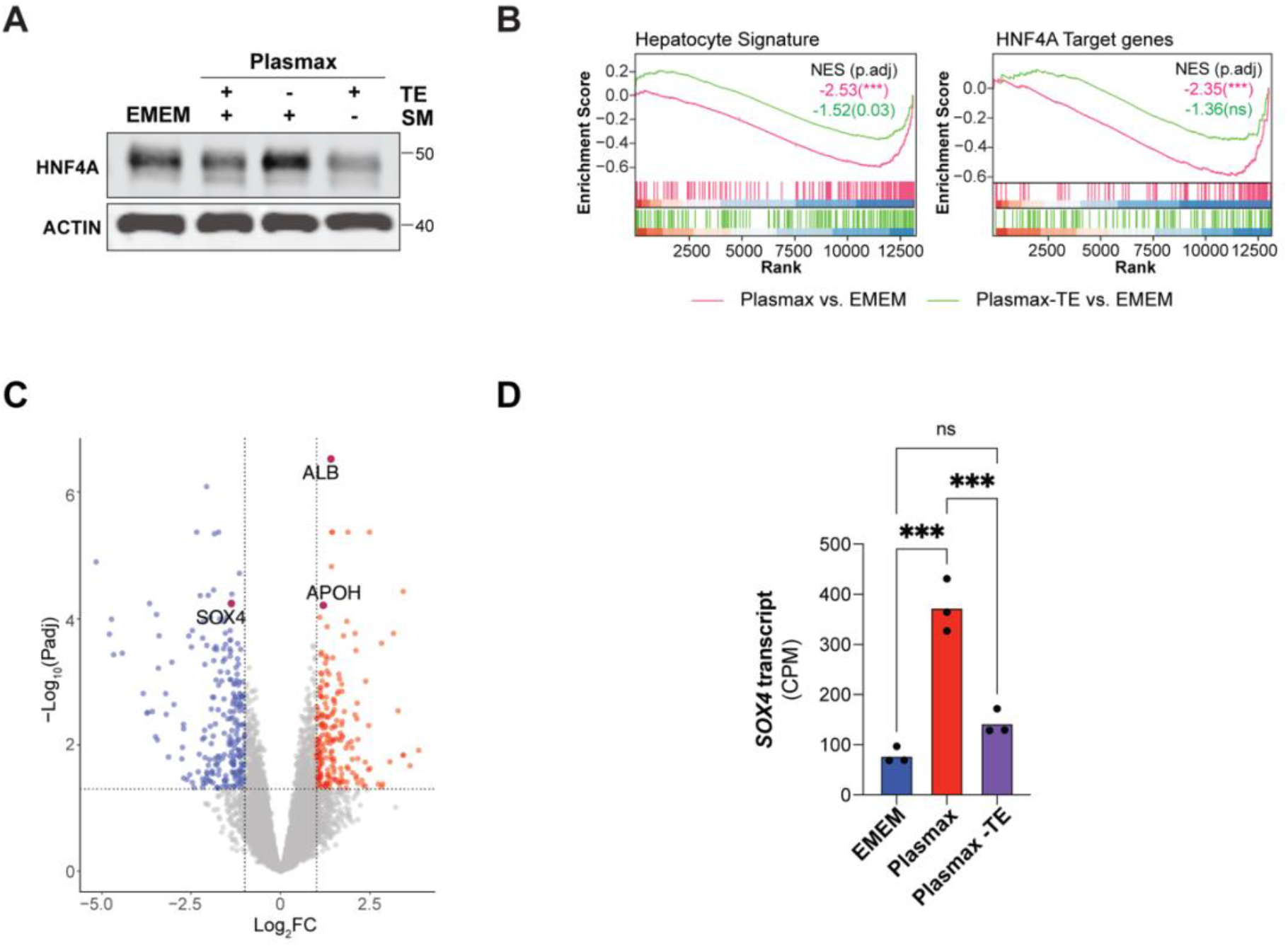
Trace element availability regulates hepatocyte differentiation. (A) Representative immunoblot analysis of HepG2 cells cultured in EMEM, Plasmax, Plasmax devoid of trace elements (TE) or Plasmax devoid of supplemental metabolites (SM). (B) GSEA plots derived from RNA-Seq analysis comparing HepG2 cells cultured in Plasmax with HepG2 cells cultured in EMEM or comparing HepG2 cells cultured in Plasmax-TE with HepG2 cells cultured in EMEM demonstrating signatures associated with hepatocyte cell state and function. (C) Volcano plot of DEGs identified from RNA-Seq analysis comparing HepG2 cells cultured in Plasmax-TE with HepG2 cells cultured in Plasmax, n=3. Hepatocyte differentiation markers are highlighted in maroon. Significantly upregulated genes are highlighted in red. Significantly downregulated genes are highlighted in blue. (D) *SOX4* transcript expression in counts per million (CPM) based on RNA-Seq analysis of HepG2 cells cultured in EMEM, Plasmax, or Plasmax-TE, n=3. For all experiments, \*\*\**P* <0.001, ns = not significant.

### Multi-omics analysis reveals a relationship between trace element availability and metalloprotein expression

To gain insights regarding which trace element(s) was contributing to dedifferentiation in Plasmax, intracellular trace element abundance was quantified using inductively-coupled plasma mass spectrometry (ICP-MS). Given that serum can be a source of trace elements, trace elements were also quantified in fresh culture media. As expected, trace elements were elevated in Plasmax compared to EMEM (Fig. S2A). Interestingly, despite iron levels being increased by only four-fold in Plasmax relative to EMEM, HepG2 cells cultured in Plasmax exhibited more than a twenty-fold increase in intracellular iron (Fig. 3A). A three-fold increase in intracellular copper was also observed (Fig. 3A). To explore the role of iron in hepatocyte differentiation, iron salts were added back to Plasmax devoid of trace elements. A multidimensional scaling (MDS) plot of the transcriptome revealed that while withdrawal of trace elements from Plasmax resulted in a transcriptome more similar to EMEM, reintroduction of iron resulted in a transcriptome that was more similar to Plasmax (Fig. 3B). This trend was recapitulated at the proteome level where reintroduction of iron restored a proteomic profile similar to HepG2 cells incubated in complete Plasmax (Fig. 3C). Interestingly, a correlation plot of the transcriptome and proteome of cells cultured in Plasmax lacking trace elements compared to complete Plasmax revealed proteins that were significantly downregulated at the protein, but not the transcript, level in response to removal of trace elements. Notably, among these proteins were selenoproteins (e.g. Selenoprotein P, SELENOP and Glutathione Peroxidase 1, GPX1) and iron-regulated proteins (e.g. Ferritin Heavy Chain 1, FTH1) (Fig. 3D). These proteins belong to the metalloprotein superfamily and their translation is dependent on trace element availability (Copeland, 2003; Addess *et al*, 1997). Consistent with this, addition of iron salts to Plasmax devoid of trace elements was able to rescue the expression of iron-regulated proteins but not selenoproteins, demonstrating the importance of trace elements in the post-transcriptional regulation of protein abundance (Fig. 3E). Taken together, these results highlight a role for trace elements in the regulation of cellular phenotype and demonstrate the importance of utilizing a multi-omic approach to interrogate the impact of altered nutrient availability.

**Figure 3.**
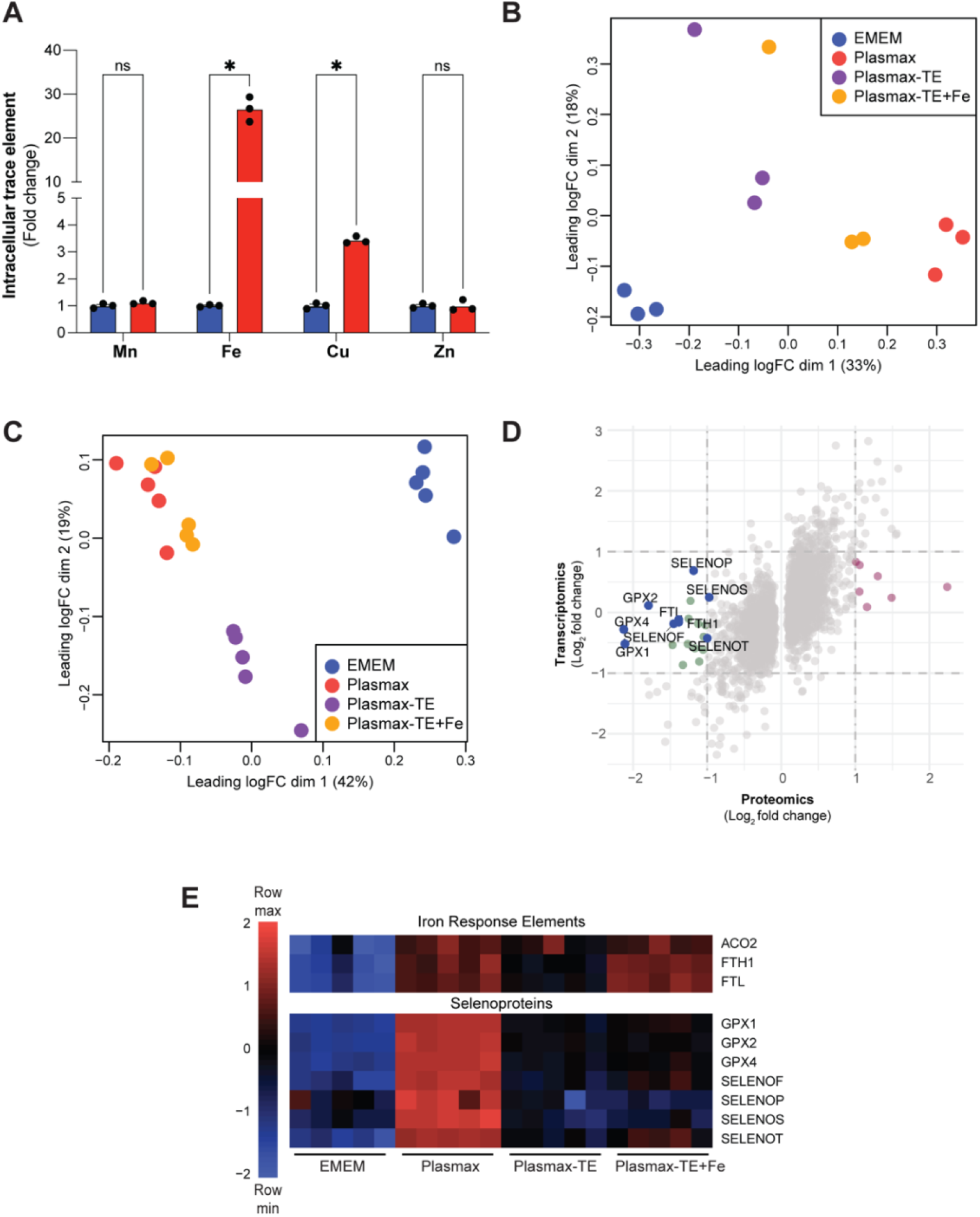
Multi-omics analysis reveals a relationship between trace element availability and metalloprotein expression. (A) Relative intracellular abundance of trace elements in HepG2 cells cultured in EMEM (blue) or Plasmax (red) n=3. (B) MDS plot of RNA-Seq data from HepG2 cells cultured in EMEM, Plasmax, Plasmax-TE, or Plasmax-TE supplemented with iron salts (0.12 µM ferric nitrate and 1.04 µM ferric sulfate; Plasmax-TE+Fe), n=3. (C) MDS plot of proteomic data from HepG2 cells cultured in EMEM, Plasmax, Plasmax-TE, or Plasmax-TE+Fe, n=5. (D) Correlation between proteomic and transcriptomic data from HepG2 cells cultured in Plasmax-TE with HepG2 cells cultured in Plasmax. Significantly differentially downregulated proteins whose transcripts were not significantly altered are highlighted in green. Significantly differentially upregulated proteins whose transcripts were not significantly altered are highlighted in magenta. Metalloproteins are highlighted in blue. (E) Heatmap of metalloprotein expression in HepG2 cells cultured in EMEM, Plasmax, Plasmax-TE or Plasmax-TE+Fe as determined by proteomic analysis, n=5. For all experiments, \**P* <0.05, ns = not significant.

### Iron availability contributes to the regulation of hepatocyte cell fate

Given that the intracellular levels of iron and copper were significantly increased in HepG2 cells cultured in Plasmax (Fig. 3A), these trace elements were further examined for their role in modulating hepatocyte cell fate. Interestingly, reintroduction of iron salts, but not copper salts, to Plasmax devoid of trace elements suppressed HNF4A expression (Fig. 4A). Moreover, re-introduction of iron salts suppressed signatures associated with the differentiated hepatocyte cell state (Fig. 4B,C). The majority of labile iron is incorporated in the cofactor heme(Ajioka *et al*, 2006). The transcriptional repressor BTB and CNC Homolog 1 (BACH1) plays a central role in regulating intracellular heme homeostasis and is itself regulated by heme availability (Oyake *et al*, 1996; Ogawa *et al*, 2001; Sun *et al*, 2002). Interestingly, BACH1 has also been shown to contribute to the regulation of cell fate (Wei *et al*, 2019; Oyake *et al*, 1996). Notably, proteomics analysis revealed that BACH1 target genes were upregulated after reintroduction of iron salts to Plasmax devoid of trace elements (Fig. 4D). Finally, BACH1 protein expression was dramatically lower in HepG2 cells cultured in Plasmax versus EMEM (Fig. 4E). Taken together, these data suggest that the iron-dependent dedifferentiation of hepatocytes in Plasmax is mediated by the trace element iron through regulation of the transcription factor BACH1 (Fig. S3A).

**Figure 4.**
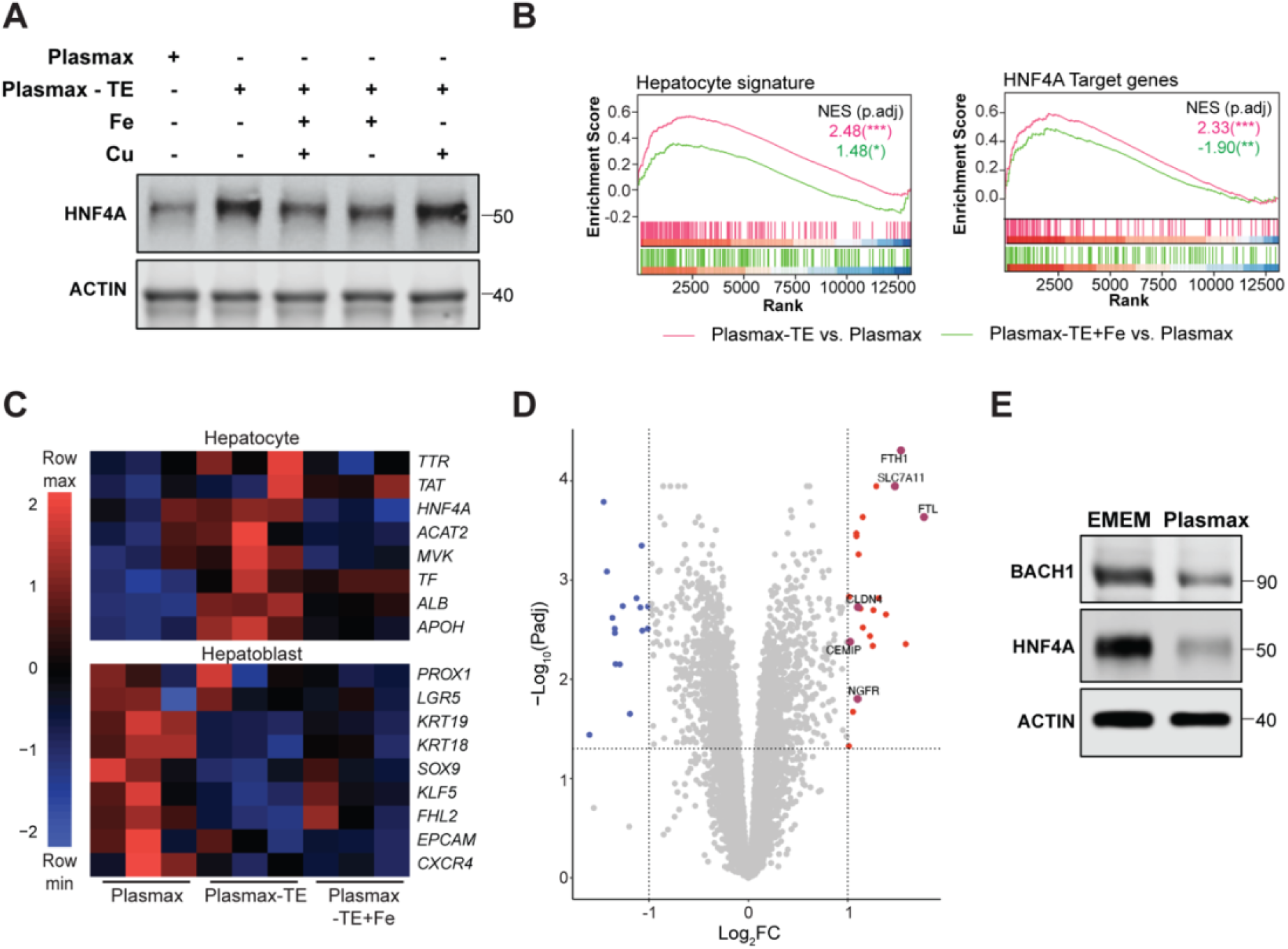
Iron availability contributes to the regulation of hepatocyte cell fate. (A) Representative immunoblot analysis of HepG2 cells cultured in Plasmax, Plasmax-TE, Plasmax-TE+Fe, Plasmax-TE supplemented with copper salts (0.005 µM cupric sulfate), or Plasmax-TE supplemented with iron and copper salts (0.12 µM ferric nitrate, 1.04 µM ferric sulfate, 0.005 µM cupric sulfate). (B) GSEA plots derived from RNA-Seq analysis comparing HepG2 cells cultured in Plasmax-TE with HepG2 cells cultured in Plasmax and comparing HepG2 cells cultured in Plasmax-TE+Fe with HepG2 cells cultured in Plasmax demonstrating signatures associated with hepatocyte cell state and function. (C) Heatmap of hepatocyte and hepatoblast gene expression in Plasmax, Plasmax-TE or Plasmax-TE+Fe as determined by RNA-Seq analysis, n=3. (D) Volcano plot of differentially expressed proteins identified from proteomic analysis comparing HepG2 cells cultured in Plasmax-TE+Fe with HepG2 cells cultured in Plasmax-TE, n=5. BACH1 targets are highlighted in maroon. Significantly upregulated proteins are highlighted in red. Significantly downregulated proteins are highlighted in blue. (E) Representative immunoblot analysis of HepG2 cells cultured in EMEM or Plasmax.

## Discussion

The liver biopsy from which HepG2 cells were isolated in 1979 was described as a well differentiated HCC (Aden *et al*, 1979). It was not until 2009 when the histopathology and genetics of the original biopsy was revisited that it was reclassified as a HB (López-Terrada *et al*, 2009). Despite this correction, HepG2 cells are still widely used to model hepatocytes/HCC rather than hepatoblasts/HB (Liu *et al*, 2022, 2021; Lin *et al*, 2016; Park *et al*, 2018; Zhang *et al*, 2018). This is reasonable as HepG2 cells cultured in conventional media exhibit characteristics consistent with differentiated hepatocytes (Fig. 1C). In this study, we have demonstrated that incubation in a physiological medium, Plasmax, causes HepG2 cells to revert to their native hepatoblast state.

A previous study investigating the impact of cell culture conditions on HepG2 cells demonstrated that increased amino acid availability promoted the acquisition of a transcriptome similar to primary human hepatocytes (Boon *et al*, 12/2020). In other cell lines, physiologically relevant metabolites that are underrepresented or absent in conventional media, such as urea and the trace element selenium, have been shown to drive significant changes in cell metabolism (Cantor *et al*, 2017; Vande Voorde *et al*, 2019). Here, we have shown that underrepresentation of the trace element iron in conventional media drives differentiation of HepG2 cells to a hepatocyte state. Maintenance of intracellular iron homeostasis has been shown to play a role in determining fate in other cell types including breast cancer cells, hematopoietic stem cells and alveolar cells (Kao *et al*, 2024; Zhuang *et al*, 2025; Müller *et al*, 2020). Although trace elements exist in minute quantities in the plasma, these studies highlight the importance of considering trace element availability when employing cell culture models.

We demonstrate that removal of trace elements from Plasmax triggers the downregulation of proteins harbouring iron response elements in the 5’-UTR of the associated transcripts (IRE; e.g. ACO2, FTL, FTH1) (Aziz & Munro, 1987; Hentze *et al*, 1987; Shen *et al*, 2023). IREs are recognized by IRE binding proteins, which in the absence of iron, repress protein expression by hindering the binding of translation initiation factors (Gray & Hentze, 1994). In addition to iron, the availability of other trace elements can dramatically impact the proteome. For example, the translation of selenoproteins, which contain the cysteine analogue selenocysteine (Sec), is dependent on selenium availability (Zinoni *et al*, 1986; Chambers *et al*, 1986; Copeland, 2003; Kryukov *et al*, 2003). Since the protein expression of IRE-containing proteins and selenoproteins are regulated at the translational level, changes in their expression resulting from altered trace element availability would not be detected at the transcriptional level. This underscores the importance of employing multi-omic approaches to investigate the consequences of altering metabolite availability.

BACH1 is a transcription factor that regulates iron homeostasis and is highly responsive to changes in iron availability (Kitamuro *et al*, 2003; Chen *et al*, 2023). Interestingly, in lung cancer, BACH1 has been shown to drive changes in cell plasticity via epithelial-to-mesenchymal transition (Wiel *et al*, 2019; Lignitto *et al*, 2019). Moreover, BACH1 has been shown to interact with transcription factors that regulate self-renewal (e.g. NANOG, SOX2, and OCT4) to repress transcription of mesodermal genes and promote pluripotency in human embryonic stem cells (Wei *et al*, 2019). Our findings demonstrate that BACH1 is downregulated in HepG2 cells cultured in Plasmax (Fig. 4E). Future studies investigating the interplay between iron availability, BACH1 and cell fate are warranted.

In conclusion, our study adds to a growing body of literature highlighting the importance of more faithfully recapitulating the *in vivo* metabolic microenvironment when employing *in vitro* cell culture models. Considering these findings, we suggest that, in addition to providing an opportunity to identify novel modulators of cell fate, physiological media will improve the *in vivo* relevance of *in vitro* cell culture models by maintaining cell fate fidelity *in vitro*.

## Materials and Methods

### Cell culture

HepG2 cells were purchased from the European Collection of Authenticated Cell Cultures (ECACC) via CellBank Australia and Hep3B cells were purchased from American Type Culture Collection (ATCC). Both cell lines were routinely assayed for mycoplasma contamination. Cells were cultured in high glucose DMEM (DMEM-HG; Thermo Fisher Scientific, 11965118), low glucose DMEM (DMEM-LG; Thermo Fisher Scientific, 11885084), Plasmax (prepared in-house as previously described (Vande Voorde *et al*, 2019)) or EMEM, which was prepared by supplementing MEM, NEAA, no glutamine (Thermo Fisher Scientific, 10370) with 1 mM sodium pyruvate, 2 mM glutamine. Cells were maintained in a humidified incubator at 37°C with 5% CO_2_. For all experiments, 2×10^5^ HepG2 cells and 8×10^4^ Hep3B cells were seeded in 6-well plates in EMEM containing 10% heat-inactivated FBS (HyClone™, SH30084). 24 hours after seeding, wells were washed with PBS and the appropriate media (DMEM-HG, DMEM-LG, Plasmax or EMEM) containing 10% heat-inactivated FBS was added. Media was refreshed after 48 hours and cells were harvested 96 hours after transition to the different media types.

### RNA-Seq analysis

For RNA-Seq analysis, total RNA was extracted using the NucleoSpin RNA kit (Macherey-Nagel, 740955) as per manufacturer’s instructions. RNA quality was confirmed using an Agilent 4200 TapeStation System. Libraries were prepared using the QuantSeq 3’ mRNA-Seq kit (Lexogen, 015) and sequenced with an Illumina NextSeq 500, with single-end 75 bp reads to a depth of 5 M reads per sample. FASTQ files were uploaded to the Galaxy web platform for quality control (FastQC), trimming (Cutadapt), alignment (RNA STAR), and counting (featureCounts) (Afgan *et al*, 2016). Reads were aligned to the human genome assembly (Ensembl hg19, GRCh37). Analysis of differentially expressed genes was performed with Limma-Voom (v3.40.6) (Smyth, 2005; Law *et al*, 2014). Gene set enrichment analysis (GSEA) was performed and plotted in R using the ‘clusterProfiler’ package (Yu *et al*, 2012). Heatmaps and volcano plots were generated in R using the ‘pheatmap’ and ‘ggplot’ packages respectively(Kolde, 2012; Wickham, 2016).

### Quantitative polymerase chain reaction (qPCR)

Total RNA was isolated using the NucleoSpin RNA Kit (Macherey-Nagel, 740955) following manufacturer’s instructions. Complementary DNA (cDNA) was synthesized using the High-Capacity cDNA Reverse Transcription Kit (Thermo Fisher Scientific, 4368813) according to the manufacturer’s guidelines. cDNA was diluted 5-fold in nuclease-free H_2_O (Thermo Fisher Scientific, 10977015). qPCR was performed using Applied Bioystems Fast SYBR^™^Green Master Mix (Thermo Fisher Scientific, 4385616) and the Applied StepOnePlus Real-Time PCR System (Thermo Fisher Scientific). Primers for *RPLP0* (5’-CAGATTGGCTACCCAACTGTT-3’ and 5’-GGGAAGGTGTAATCCGTCTCC-3’), *ALB* (5’-CTTGAATGTGCTGATGACAGG-3’ and 5’-GCAAGTCAGCAGGCATCTCAT-3’), and *ACAT2* (5’-GCGGACCATCATAGGTTCCTT-3’ and 5’- CTGCAAGGCACACAGCTTTT-3’) were used for qPCR. Relative gene expression was determined using the ΔΔCt method(Rao *et al*, 2013) normalised to *RPLP0*.

### Immunoblotting

Cells were washed with PBS and lysed in SDS lysis buffer (1% SDS, 50 mM Tris-HCl, 10 mM EDTA) containing a protease inhibitor cocktail (Sigma-Aldrich, P8340) and Pierce™ Universal Nuclease for Cell Lysis (Thermo Fisher Scientific, 88072). Lysates were resolved by SDS-PAGE and transferred to nitrocellulose membrane. Membranes were probed with primary antibodies recognising HNF4A (Cell Signaling Technology, 3113, 1:1000 dilution), BACH1 (Proteintech, 14018-1-AP, 1:500 dilution), and β-Actin (Cell Signalling Technology, 3700, 1:5000 dilution). Membranes were incubated with IRDye secondary antibodies (LI-COR, 926- 68070 and 926-32211) and imaged using the Odyssey DLx imaging system (LI-COR).

### Inductively-coupled plasma mass spectrometry (ICP-MS)

ICP-MS was performed as previously described(Pyun *et al*, 2022). Cells were washed with PBS and whole cell lysates were collected in SDS lysis buffer. Media with 10% FBS (500 µL) was used to determine trace element abundance in EMEM and Plasmax. Lysates or media were lyophilized prior to digestion in 65% nitric acid (Merck, 100441). Samples were heated at 90^°^C for 20 minutes to complete the digestion and subsequently diluted with 1% (v/v) nitric acid. Samples were run on the Agilent 8800 (ICPMS-QQQ-8800) system using a Helium reaction gas cell. Raw parts per billion (ppb) data of the elements were converted to μmol/L with dilution factors applied. The final concentrations of the metals were corrected with the treatment blanks and normalized to the total protein to obtain a final unit expressed as metal/mg protein. Fold changes were normalized to the average of controls.

## Proteomic Analysis

Cells were washed with PBS and lysed in 200 µL of ice-cold RIPA buffer (1% NP-40, 0.5% sodium deoxycholate, 0.1% SDS, 150 mM NaCl, 50 mM Tris-HCl, pH 7.5) containing a protease-inhibitor cocktail (Sigma-Aldrich, P8340) and phosphatase inhibitors (Roche, 4906845001). Protein (50 µg per sample) was reduced by adding a final concentration of 10 mM TCEP (Sigma-Aldrich, 646547) and incubated at 37^°^C for 45 min. Samples were then alkylated with 50 mM iodoacetamide (Sigma-Aldrich, I1149) and incubated in the dark for 45 min. After acidification with 2.5% phosphoric acid (Sigma-Aldrich, 79617), samples were passed through S-Trap micro columns (Protifi, C02-micro). Proteins were digested overnight at 37^°^C with a 1:20 dilution of Pierce™ Trypsin Protease (Thermo Fisher Scientific, 90057). Peptides were eluted sequentially with 50 mM triethylammonium bicarbonate buffer (TEAB), 0.2% formic acid, and 50% acetonitrile/0.2% formic acid. Finally, the eluent was lyophilized and reconstituted in 2% acetonitrile/0.05% trifluoroacetic acid to make up the peptide solution for LC-MS/MS analysis by data-independent acquisition using a Thermo OrbiTrap Ascend mass spectrometer. Data analysis was performed using the software suite, DIA-NN (Demichev *et al*, 2020). Heatmaps and volcano plots were generated in R using the ‘pheatmap’ and ‘ggplot’ packages respectively (Kolde, 2012; Wickham, 2016).

## Quantification and Statistical Analysis

Statistical analyses were performed with Prism 9 software (GraphPad Software). All statistical analyses for data comparing two groups were performed with an unpaired Student’s t-test. One-way ANOVA with the Holm–Sidak method for multiple comparisons was used for comparison of more than two groups. All immunoblots are representative of results from at least three independent experiments. All other statistical details of experiments can be found in the figure legends.

## Acknowledgements

A.J.S.O. is supported by a Peter MacCallum Cancer Centre Foundation Grant. T.A.T. is supported by an Australian Government Research Training Program Scholarship. K.K.B. is supported by NHMRC Ideas Grants (GNT2004212 and GNT2012313) and a Victorian Cancer Agency Mid-Career Research Fellowship (MCRF17020). A.G.C. is supported by a National Health and Medical Research Council (NHMRC) Investigator Grant (GNT1176650), and an Australian Research Council Discovery Project Grant (DP200102693). K.K.B and A.G.C. are also jointly supported by the Peter MacCallum Cancer Foundation (Ted and Lila Seehusen Foundation). We acknowledge support from the Peter MacCallum Cancer Centre Foundation and the Australian Cancer Research Foundation. We acknowledge the Bio21 Melbourne Mass Spectrometry and Proteomics Facility (MMSPF) at the University of Melbourne and the Biometals Facility at the Florey Institute of Neuroscience and Mental Health (University of Melbourne) for their training, support, and technical assistance. We extend our thanks to the Peter MacCallum Cancer Centre Core Facilities and their staff who provided support for this work; namely the Molecular Genomics Core, the Flow Cytometry Core and the Bioinformatics Core Facilities. Finally, we thank members of the Cox Laboratory and Brown Laboratory (Peter MacCallum Cancer Centre) for helpful discussions.

## Author contributions

A.J.S.O., A.G.C., and K.K.B. designed research; A.J.S.O., T.A.T., and J.M.R. performed research; A.J.S.O. analyzed data; A.G.C. and K.K.B. supervision; and A.J.S.O., A.G.C., and K.K.B. wrote the paper.

