## Supplementary Information for "A multi-omic approach reveals iron availability influences cell fate fidelity"

**Ong et al.**

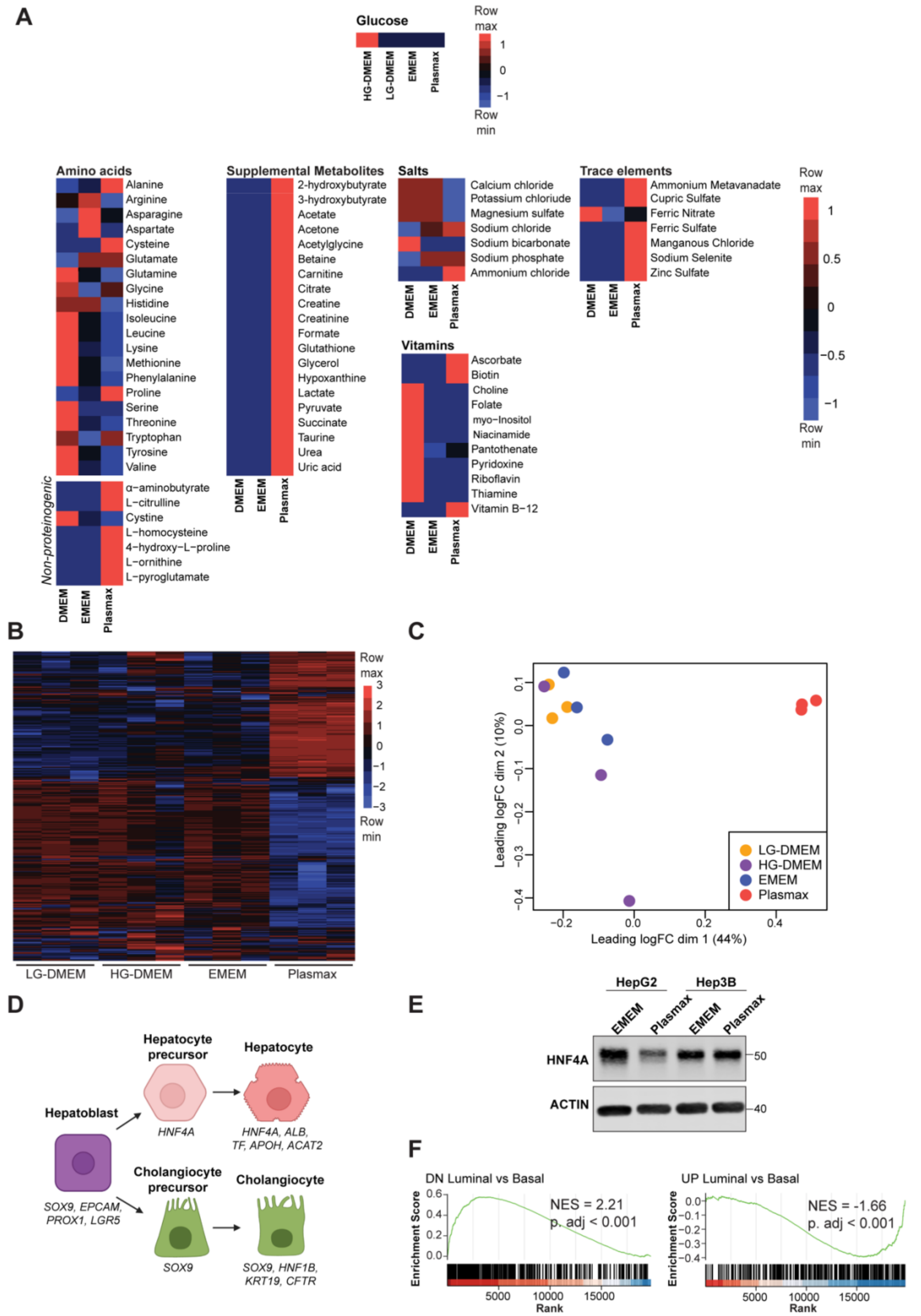


**Figure S1. HepG2 cells adopt a native hepatoblast-like state in physiological media.** (A) Heatmap of the relative concentrations of the components of DMEM (high glucose, HG; low glucose, LG), EMEM, and Plasmax. (B) Heatmap of the top 500 variable genes expressed in LG-DMEM, HG-DMEM, EMEM or Plasmax as determined by RNA-Seq analysis, n=3. (C) MDS plot of RNA-Seq data from HepG2 cells cultured in LG-DMEM, HG-DMEM, EMEM or Plasmax, n=3. (D) Schematic of liver cell differentiation during development. (E) Representative immunoblot analysis of HepG2 and Hep3B cells cultured in EMEM or Plasmax. (F) GSEA plots derived from RNA-Seq analysis comparing MDA-MB-468 cultured in Plasmax with MDA-MB-468 cells cultured in DMEM-F12 demonstrating signatures associated with breast cancer cell state.

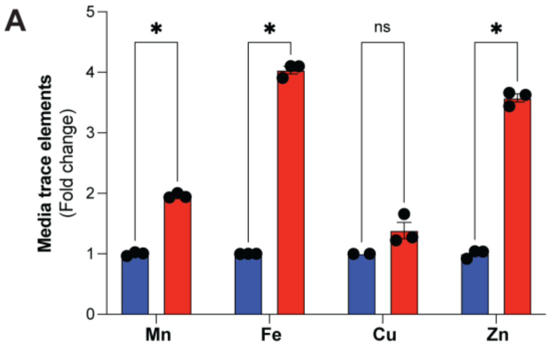


**Figure S2. Multi-omics analysis reveals a relationship between trace element availability and metalloprotein expression**

(A) Quantification of the relative abundance of trace elements in EMEM (blue) or Plasmax (red), n=3. For all experiments, **P* <0.05, ns = not significant.

A

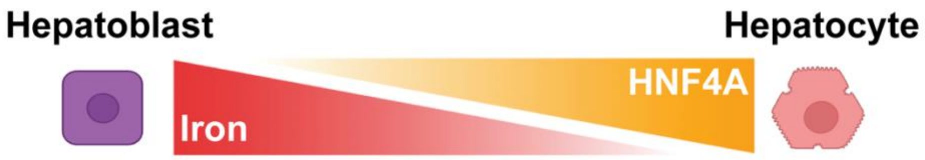


**Figure S3. Iron availability contributes to the regulation of hepatocyte cell fate.**
(A) Schematic demonstrating the reciprocal relationship between iron availability, HNF4A activity and hepatic fate.
